# QuicK-mer: A rapid paralog sensitive CNV detection pipeline

**DOI:** 10.1101/028225

**Authors:** Feichen Shen, Jeffrey Kidd

## Abstract

QuicK-mer is a unified pipeline for estimating genome copy-number from high-throughput Illumina sequencing data. QuicK-mer utilizes the Jellyfish application to efficiently tabulate counts of predefined sets of k-mers. The program performs GC-normalization using defined control regions and reports paralog-specific estimates of copy-number suitable for downstream analysis. The package is freely available at https://github.com/KiddLab/QuicK-mer

## 1 Introduction

Detecting copy-number variation (CNV) from high-throughput sequencing data is a prevalent and important problem in the research of genome evolution, population genetics, and disease. Several methodologies have been developed that make use of distinct features of the data – including read depth, read-pair mappings, split-reads, sequence assembly, and combinations of multiple signals (Alkan *et al.*, 2011). However, current methods are not without limitations. For example, read depth approaches rely upon mapping to a reference genome assembly and are typically unable to isolate variation among duplicated sequences present in multiple locations in the reference. Specialized approaches have been developed to address this limitation, including tabulating depth at paralog-specific nucleotide positions using customized tools (Sudmant *et al.*, 2010; Alkan *et al.*, 2009; Handsaker *et al.*, 2015). Although effective, these approaches require re-analysis of existing data using specialized mapping programs, such as mrsFAST (Hach *et al.*, 2010), or extensive downstream processing both of which can add considerable time to the analysis. Additionally, previous approaches require trimming and/or partitioning of sequence reads into subsequences of a specified length and therefore do not make full use of the available sequence data. In this application note, we present QuicK-mer, a rapid and paralog-sensitive CNV estimation pipeline that efficiently produces copy-number estimates from FASTQ or BAM input files in less than 6 hours. Our approach is mapping-free and relies upon efficient tabulation of read depth at predefined sets of informative k-mers.

## 2 Methods

To achieve efficient and paralog-specific CNV estimation, we focused on counting specific k-mer sequences rather than aligning reads to a reference, an approach that has also been proposed for analysis of RNA-Seq data (Zhang and Wang, 2014; Patro *et al.*, 2014). The QuicK-mer pipeline is designed to utilize the existing Jellyfish k-mer counting application (Marçais and Kingsford, 2011). Accepting both FASTQ and BAM files as input, QuicK-mer is designed for sequences generated by the Illumina platform.

The QuicK-mer pipeline requires several pre-processing steps that must occur once for each species reference genome utilized. First, a predefined set of informative k-mers must be identified. For genome-wide CNV analysis, we utilize a catalog of unique 30-mers that do not overlap with highly repeated sequences (Supplementary Methods). The k-mer list is sorted based on coordinates in the reference assembly and paired with two additional files: a file describing the GC content within 400 bp of each k-mer and a file containing a binary flag indicating whether each k-mer should be considered as a control utilized for copy-number normalization (Supplementary Methods). Each of these two files is stored in an efficient binary format. Precomputed files for recent human, mouse, and dog reference genomes are publicly available and generation procedures are further described in the Supplementary Methods.

For CNV estimation, the QuicK-mer pipeline uses Jellyfish-2 to tabulate k-mer counts for each k-mer in the target catalog using the sequencing file input. If a BAM file is specified by the user, QuicK-mer will optionally filter reads based on the duplicate flag using Samtools (Li *et al.*, 2009). To increase the efficiency of k-mer counting, we initialize a bloom counter in Jellyfish-2 with two copies of the k-mer catalog. This ensures that only the k-mers of interest are tabulated, thus reducing memory and disk I/O burden. Next, the counts for each k-mer are extracted from the Jellyfish database and stored in memory. Each k-mer is then checked for status as a normalization control and, if indicated, incorporated into the construction of a GC-bias correction curve. To calculate GC bias, we average the depth for all the k-mers having the same 400bp GC percentage in increments of 0.25%. A correction is then calculated to normalize the average depths in all GC bins, which is then applied to all interrogated k-mers. The resulting normalized k-mer counts can then be converted to copy-number estimates based on the assumed copynumber of indicated control regions (CN=2 for autosomes, see Supplementary Methods). The resulting copy-number estimates are then exported at the level of individual k-mers or in defined sets of windows for subsequent analysis and visualization. Standard segmentation algorithms can be employed on the QuicK-mer output to identify copy-number change points within an individual genome. Alternatively, copy-number estimates for specific intervals can be compared across samples to define locus-specific copy-number genotypes.

## 3 Results

We have tested QuicK-mer using Illumina short read WGS data with various levels of sequencing coverage (1-23x, Supplementary Table 1). The majority of data was obtained from the 1000 Genome Phase 1 and 3 and pilot data, as well as higher-coverage datasets from a previously analyzed set of samples (Bentley *et al.*, 2008; Abecasis *et al.*, 2010, 2012; Auton *et al.*, 2015). Efficiency assessments demonstrate that QuicK-mer is able to estimate genome-wide copy number for a human genome sequenced to 17x in less than 6 hours using 4 cores on a typical computing cluster.

To account for the bias caused by the local GC content in the genome, we ran QuicK-mer on sequencing lanes individually. Supplementary Figure 2 shows the distribution of GC curves observed from different sequencing lanes within the same WGS shotgun library. Once the corrected depth is obtained, all files from the same sample were merged. For comparison, we set the display resolution to be 500 30-mers per window and reported the median copy number. We compared with the supplementary data reported in Sudmant *et al*., 2010 and find that QuicK-mer can distinguish paralog-specific copy number in duplicated regions of the human genome (Fig 1 and Supplementary Figures 5-14). Thus, QuicK-mer is a map-free approach that scales well, requires minimal additional processing, and represents a unified pipeline for efficiently estimating paralog-specific copy-number from Illumina WGS data.

**Fig 1.**
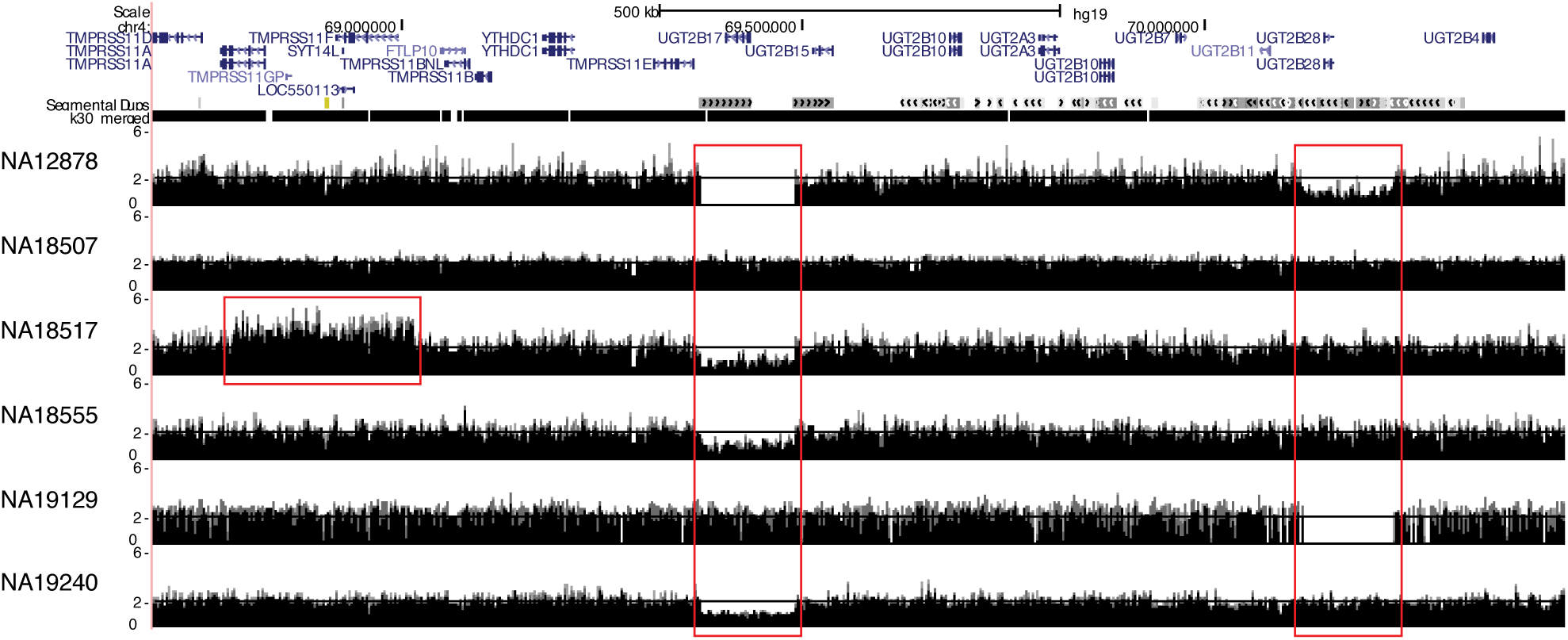
Diverse CNV detected by QuicK-mer. Paralog-specific copy numbers are estimated in the chr4q13.2 *UBT2* gene family region. Red boxes indicate regions of copy-number variation. *UGT2B17* is hemizygously deleted in NA19240, NA18555 and NA18517. *TMPRSS11F* and *SYT14L* are duplicated in NA18517 resulting in a copy number of 3. This figure corresponds to the region shown in Figure S65 in Sudmant *et al.*, 2010. The k30_merged track indicates the locations with unique 30-mers.

## Acknowledgements

We thank Ryan Mills for comments on manuscript drafts and Lukas Kuderna and Tomas Marques-Bonet for evaluating the QuicK-mer pipeline.

## References

Abecasis, G.R. et al. (2012) An integrated map of genetic variation from 1,092 human genomes. Nature, 491, 56–65.

Abecasis, G.R.R. et al. (2010) A map of human genome variation from population-scale sequencing. Nature, 467, 1061–73.

Alkan, C. et al. (2011) Genome structural variation discovery and genotyping. Nat. Rev. Genet., 12, 363–76.

Alkan, C. et al. (2009) Personalized copy number and segmental duplication maps using next-generation sequencing. Nat. Genet., 41, 1061–7.

Auton, A. et al. (2015) A global reference for human genetic variation. Nature, 526, 68–74.

Bentley, D.R. et al. (2008) Accurate whole human genome sequencing using reversible terminator chemistry. Nature, 456, 53–59.

Hach, F. et al. (2010) mrsFAST: a cache-oblivious algorithm for short-read mapping. Nature methods, 7, 576–577.

Handsaker, R.E. et al. (2015) Large multiallelic copy number variations in humans. Nat. Genet., 47, 296–303.

Li, H. et al. (2009) The Sequence Alignment/Map format and SAMtools. Bioinformatics, 25, 2078–9.

Marçais, G. and Kingsford, C. (2011) A fast, lock-free approach for efficient parallel counting of occurrences of k-mers. Bioinformatics, 27, 764–70.

Patro, R. et al. (2014) Sailfish enables alignment-free isoform quantification from RNA-seq reads using lightweight algorithms. Nature biotechnology, 32, 462–464.

Sudmant, P. et al. (2010) Diversity of Human Copy Number Variation and Multicopy Genes. Science, 330, 641–646.

Zhang, Z. and Wang, W. (2014) RNA-Skim: a rapid method for RNA-Seq quantification at transcript level. Bioinformatics, 30, i283–i292.

